# Protein phosphatase V ensures timely cell cycle remodeling during the mid-blastula transition in Drosophila

**DOI:** 10.1101/172379

**Authors:** Boyang Liu, Hung-wei Sung, Ingo Gregor, H.-Arno Müller, Jörg Großhans

## Abstract

Cell cycle remodeling from fast nuclear cycles to a generic cell cycle mode is a major feature of the mid-blastula transition (MBT) in *Drosophila.* Remodeling occurs when Twine/Cdc25 falls below a critical threshold. Timing is based on Twine destabilization induced by zygotic transcription. It is conceivable that appropriate starting levels are also important for timely reaching the threshold. Mechanisms for controlling Twine levels at the onset of MBT are unknown. Here we identify a function of the protein phosphatase V in this mechanism. Twine was increased in *P_p_V* mutants, whereas the decay rate was comparable to wildtype. *P_p_V* mutants frequently underwent an extra nuclear division. We detected P_p_V-dependent phosphosites in Twine. Phosphosite mutants contain higher Twine levels and frequently underwent an extra nuclear division, comparable to *P_p_V* mutants. Our data support a model that the cell cycle remodeling is controlled by induced destabilization and P_p_V-d ependent control of Twine levels.

## Introduction

A change in the mode of the cell cycle from a fast nuclear cycles with no gap phases and no cytokinesis to a generic mode with a long gap phase and slow replication is a characteristic feature of the mid-blastula transition (MBT) in early embryogenesis (Farrell 2014, Blythe2015a, Liu2017). This remodeling of the cell cycle is linked to changes in a number of cellular processes, including DNA replication checkpoint, epigenetic markers, onset of zygotic gene expression, degradation of maternal RNAs and morphological changes (Blythe 2015a, Harrison2015, Laver 2015). In *Drosophila* embryos, the cell cycle remodeling occurs after the mitosis of nuclear cycle 13 (Foe1993). The activation of zygotic transcription triggers the cell cycle switch in that the DNA checkpoint is activated (Sung2013, Blythe2015b) and few zygotic genes encoding mitotic inhibitors are expressed (Grosshans 2000, Grosshans 2003, Gawlinski2007).

At the center of the embryonic cell cycle switch is a positive regulator of Cdk1, the phosphatase Twin e/Cdc25 (Edgar 1996), which is stable during the 13 fast cycles but is destabilized by zygotic factors like Tribbles (Trbl) after mitosis 13 (Di Talia2013, Farrell 2013). Twine levels normally fall below the critical threshold for entry into mitosis in interphase 14, which is achieved by a drop in half life by a factor of 4 from about 20 min to 5 min (Di Talia2013). Since there is significant protein synthesis, the decay time of Twine, including translation and degradation, is longer than its half life. However, important for when the levels of Twine reach the threshold is not only the rate of decay. It is conceivable that an increase in the starting levels leads to a delay in reaching the threshold. Mechanism for controlling Twine protein expression at the onset of MBT are unknown.

The *Drosophila Protein phosphatase V* (*P_p_V*) encodes the homologue of the catalytic subunit of human PP-6 (Mann1993, Bastians1996). PP-6 has been implicated in mitosis and meiosis (Stefansson2007, Chen2007, Hu2015, Ertych2016), DNA repair (Zhong2011), inflammation (Yan2015), *C. elegans* cortical contractility (Afshar2010) and mouse early embryogenesis (Ogoh2016). In addition to these physiological functions, PP-6 mutations have been found to be associated with melanoma tumors (Hodis 2012, Krauthammer 2012, Hammond 2013). PP-6 is an inhibitory phosphatase of oncogene AuroraA kinase (Zeng2010, Ertych2016). Quantitative phosphoproteomics identified further potential PP-6 substrates (Rusin 2015), whose physiological relevance has remained undefined.

Here we characterize the function of *P_p_V* in cell cycle remodeling during MBT in *Drosophila* embryos. We isolated and characterized a *P_p_V* null mutant allele. We reveal a function in controlling the switch in cell cycle mode. Loss of *P_p_V* frequently led to an additional nuclear division and increased the variance of Twine protein levels. Our results indicate that P_p_V controls low initial levels of Twine, allowing a timely drop of protein levels below the threshold for entry into mitosis.

## Results

### *P_p_V* mutant embryos frequently undergo an extra nuclear division cycle

We screened a collection of mutations with early embryonic phenotypes (Luschnig2004, Vogt2006) for cell cycle defects (Fig. 1A). Embryos from females with germline clones of the X9 mutation (in the following designated as *P_p_V*[X9] embryos) developed apparently normally until gastrulation but underwent frequently one extra round of nuclear division as shown by time lapse imaging with bright field microscopy and with embryos with fluorescently labelled histone (Fig. 1B, Movie 1). The extra division often involved only part of the embryo resulting in regions with higher nuclear density (Fig. 1B). The penetrance of the extra division (including divisions involving only a part of the embryos) was in the range of one third to one half of the embryos. The timing of the early cell cycles was comparable to wild type embryos (Fig. 1C). Cycle 13 was slightly prolonged in *P_p_V*[X9] embryos with a normal number of nuclear divisions and similar to the length of cycle 14 in *P_p_V*[X9] embryos with an extra nuclear division. Many eggs from *P_p_V*[X9] germ line clones (about 50%, N>100) did not develop, indicating a function in egg activation, meiosis or fertilization.

**Figure 1:**
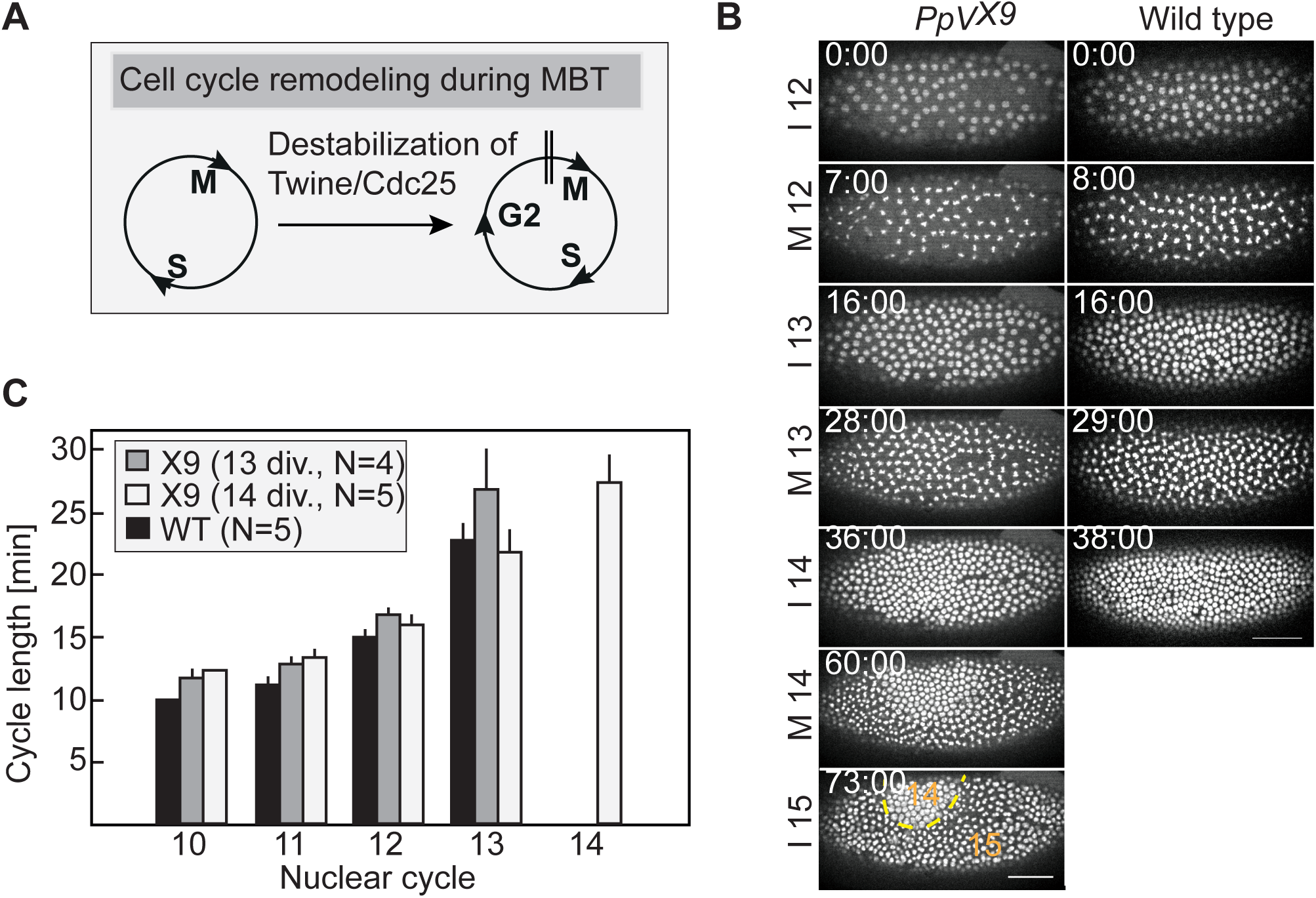
P_p_V mutants frequently undergo 14 nuclear divisions. **(A)** Scheme for the cell cycle remodeling during MBT. **(B)** Images from time lapse recordings of wild type embryos and embryos from *P_p_V*[X9] germ line clones expressing Histone 2Av-RFP and their **(C)** quantification for cell cycle lengths. *P_p_V*[X9] embryos were grouped in ones with normal cell cycle number (13 divisions) and ones with at least a partial extra cycle (14 divisions). Bars indicate mean values, associated lines indicate standard error of the mean. Scale bar 50 μm.

We mapped the lethality and the cell cycle phenotype by meiotic recombination and complementation with duplication and deficiency chromosomes to the 5F region (Supplemental material Fig. S1). Sequencing of the genes in the mapped region revealed a nonsense mutation in the seventh codon in the *P_p_V* gene encoding the *Drosophila* homologue of the catalytic subunit of protein phosphatase 6 (Fig. 2A, 2B). This mutation is responsible for the phenotype, since a genomic construct rescued the lethality and the cell cycle phenotype of *P_p_V*[X9] (Fig. 2A). *P_p_V*[X9] is likely to be a protein null allele as a band was detected by western blot with extracts from wild type and rescued but not *P_p_V* mutant embryos (Fig. 2C). We did not detect a band in extracts from staged embryos (Fig. 2C), indicating a lack of significant zygotic expression. Consistently, we did not observe a zygotic rescue leading to an ameliorated phenotype or larval hatching in half of the embryos.

**Figure 2.**
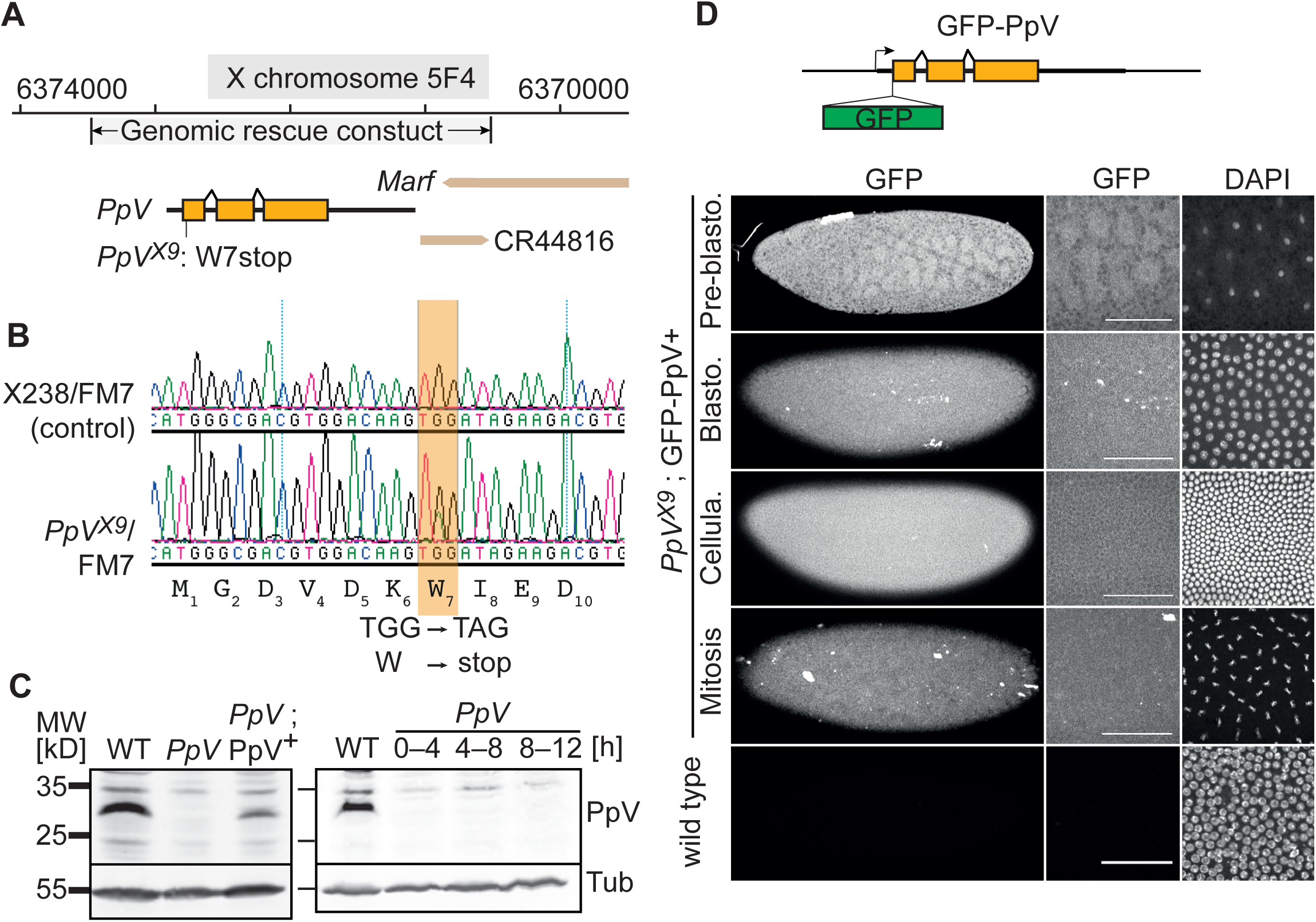
Identification of a *P_p_V*[X9] mutation. **(A)** Map of the *P_p_V* locus. The X9 mutation leading to a stop codon at position 7 is indicated (W7 stop). **(B)** Sequence traces of DNA amplified from heterozygous *P_p_V*[X9]/FM7 and in comparison X238/FM7 with a related genetic background. The double peak in the X9 trace reveals a non-sense mutation in codon 7. **(C)** Western blot with extracts (0–4 h and as indicated) of wild type embryos and embryos from germ line clones of *P_p_V*[X9] and *P_p_V*[X9]; P_p_V[+] crossed with wild type males. Loading control with α-Tubulin (Tub). **(D)** Fixed embryos with and without a genomic rescue construct GFP-P_p_V[+] were stained for GFP and DAPI. Scale bar 50 μm.

Due to cross reactivity, the antiserum was not suitable for immunostaining. Employing a transgene with a GFP inserted at the N-terminus of P_p_V, which rescued lethality and the embryonic cell cycle phenotype, we could specifically detect the distribution of P_p_V protein (Fig. 2D). We detected a spatially and temporally uniform expression of GFP-P_p_V in early embryos. Close observation revealed slightly lower nuclear levels. We did not observe obvious differences in fluorescence among embryos in pre-gastrulation stages and embryos in interphase and mitosis, suggesting that expression levels of P_p_V protein did not change during the early embryonic cell cycles (Fig. 2D).

### Twine protein levels are increased and longer persisting in *P_p_V* mutants

Cell cycle remodeling at MBT is induced by destabilization of Twine/Cdc25 (Di Talia2013, Farrell2013). We investigated genetic interactions of *P_p_V* and *twine*. We found that *twine* is a dominant suppressor of *P_p_V*, since we observed a few hatching larvae from *P_p_V* germ line clones that were heterozygous for *twine* (5%, N>1000). This suggests that *P_p_V* may act in an antagonistic manner on *twine*. Total *twine* RNA as measured by qPCR was similar in wild type and *P_p_V* embryos (1: 0.94). We compared the developmental profile of Twine protein in wild type and *P_p_V* embryos by immunostaining of fixed embryos for Twine. We found that Twine staining tentatively persisted longer in *P_p_V* embryos than wild type embryos as staged by morphological markers (Fig. 3). However, comparison of Twine levels between wild type and mutant embryos is problematic, since Twine levels were highly dynamic in cycle 14 and showed high variance especially in the *P_p_V* mutants. For reliable and precise quantification of the rapid and variant dynamics, a noninvasive quantitative method is needed that is applicable to single embryos during MBT.

**Figure 3.**
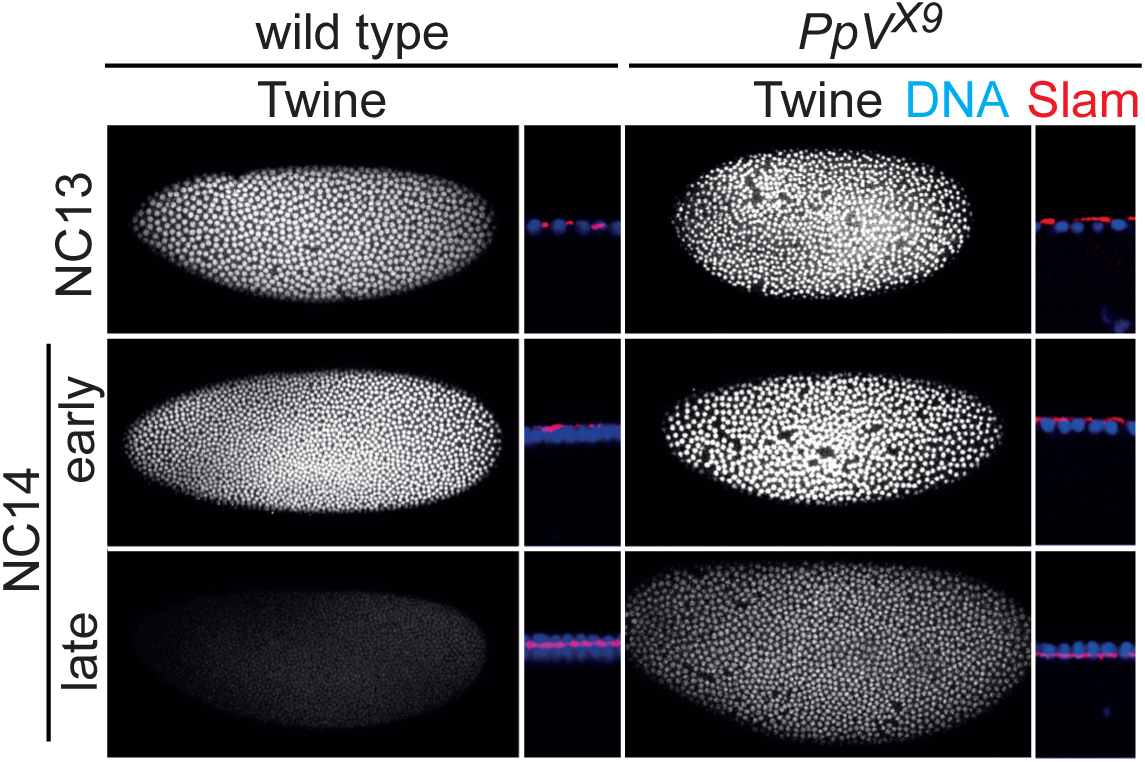
P_p_V modulates Twine protein expression. Fixed wild type embryos and embryos from germ line clones of *P_p_V*[X9] stained for Twine (white), Slam (red) and DNA (blue). Sagittal sections with Slam staining and nuclear morphology allows approximate staging. NC: nuclear cycle.

We applied fluctuation analysis employing embryos expressing Twine-GFP protein in order to reveal the temporal profile in individual embryos (Fig. 4A–C). As the fluctuations in the fluorescence traces are dependent on the GFP concentration, the number of molecules per optical volume can be calculated (Chen2002, Digman2008, Fig. 4C). We validated the method in embryos expressing one or two copies of nuclear localization signal GFP (nls-GFP) driven by the ubiquitin promoter. Western blotting with extracts from these embryos demonstrated that protein levels corresponded to gene copy number (Fig. 4D). Fluctuation analysis with embryos in early cycle 14 revealed an increase from c(1× GFP) = 102.3±17.3 to c(2×GFP)=213.7±49.1, and thus a two-fold higher absolute concentration of nls-GFP in embryos with two transgenes as compared to embryos with one nls-GFP transgene (Fig. 4E).

**Figure 4.**
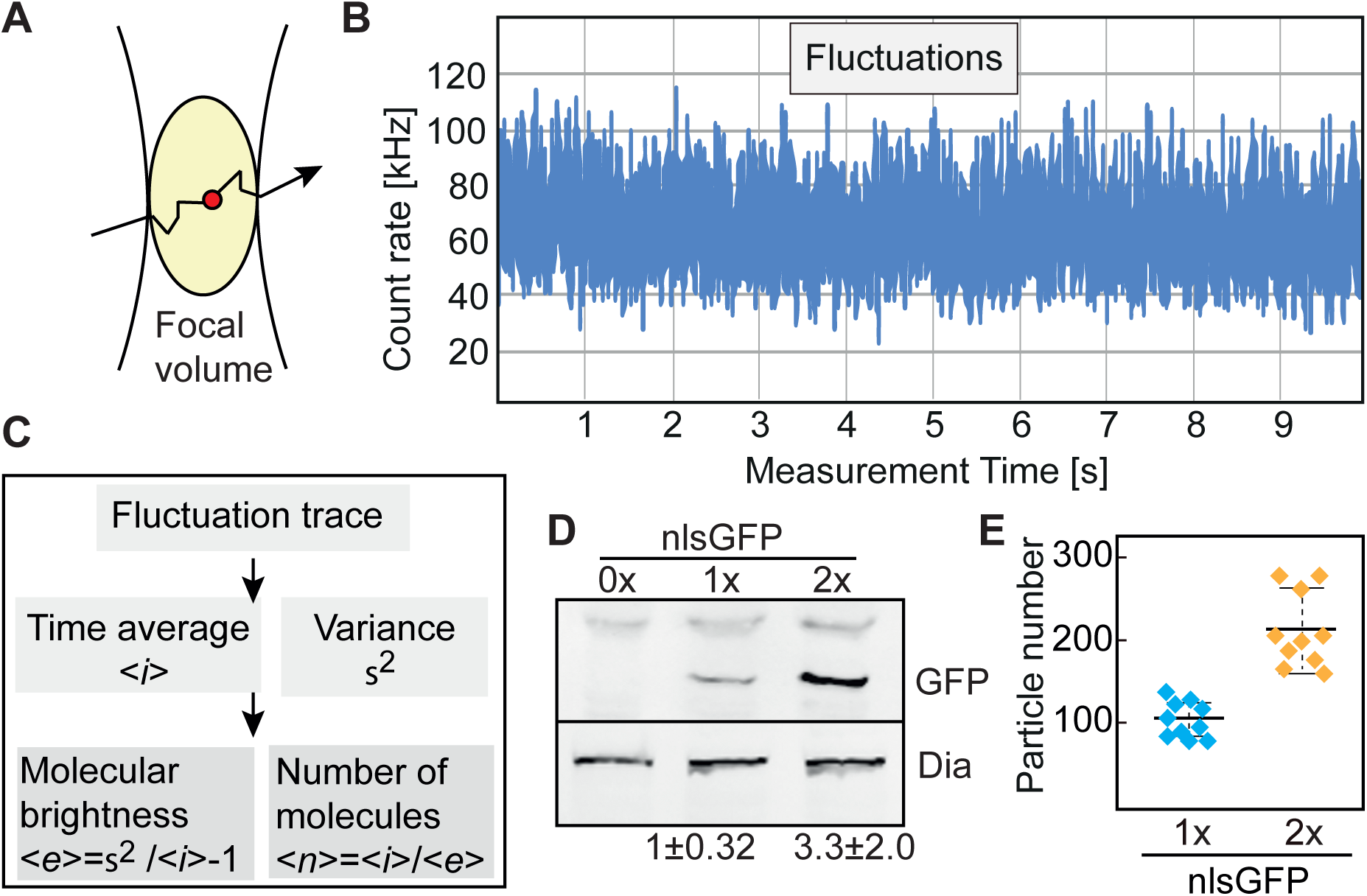
Fluctuation analysis for absolute concentration dynamics. **(A)** Principle of fluctuation analysis. Passing through the focal volume leads to changes in the fluorescence signal. The frequency of these changes dependent on the concentration. **(B)** Fluctuation trace of a measurement. **(C)** Time average and variance are computed from the fluctuations trace, which allows calculation of average molecular brightness and average number of molecules within the focal volume. **(D)** Western blot with extracts of the embryos (0–3 h) with 0×, 1×, 2× copies of an nls-GFP transgene. Quantification by densitometry (N = 3). **(E)** Particle number per focal volume as determined by fluctuation analysis of GFP fluorescence. Mean, bold line; standard deviation, dashed line.

Having validated the method, we recorded the concentration profile of Twine-GFP fluorescence during the course of interphase 14 (Fig. 5B). The Twine-GFP transgene is fully functional in *twine* mutants as female sterility was rescued (Fig. 5A). From the time courses of Twine-GFP concentration in single embryos (Fig. 5C), we calculated two parameters for each embryo assuming an exponential function, the decay time (which includes protein production and degradation and is larger than the half life of the protein) and the initial particle number, which corresponds to the concentration at the onset of interphase 14 (Fig. 5C, 5D). In embryos with one copy of Twine-GFP in addition to the two endogenous copies of *twine*, we calculated values with little variation around the mean (12.9%, Tab. 2), indicating that our method is robust (Fig. 5E, 5F). In contrast, a statistically significant change of the initial particle number from 38.5±5.0 in wild type to 56.7 + 19.5 in *P_p_V* mutants was calculated (Tab. 2). More striking than the change of the mean value is the strongly increased variance. The standard deviation more than doubled from 13% in wild type to 34% in *P_p_V* mutants (Fig. 5E, Tab. 2). Many of the *P_p_V* mutants have an initial partial number of approximately 80 that is double of the wild type number while others are in the range of wild type. The decay time of about 10 min remained unchanged in *P_p_V* mutant embryos (Fig. 5F). Visa versa, we found a statistically significant change in the decay time to about 13.5 min in *trbl* embryos (Fig. 5F, Tab. 2) but an unchanged initial particle number (Fig. 5E). These data are consistent with the previous report on the role of *trbl* in Twine destabilization (Farrell 2013). These measurements indicate that P_p_V controls the expression levels of Twine protein and maintains a low embryo to embryo variation in pre-MBT. Complementary to P_p_V, zygotic Trbl is involved in the induced destabilization of Twine during MBT by enhancing the decay rate.

**Figure 5.**
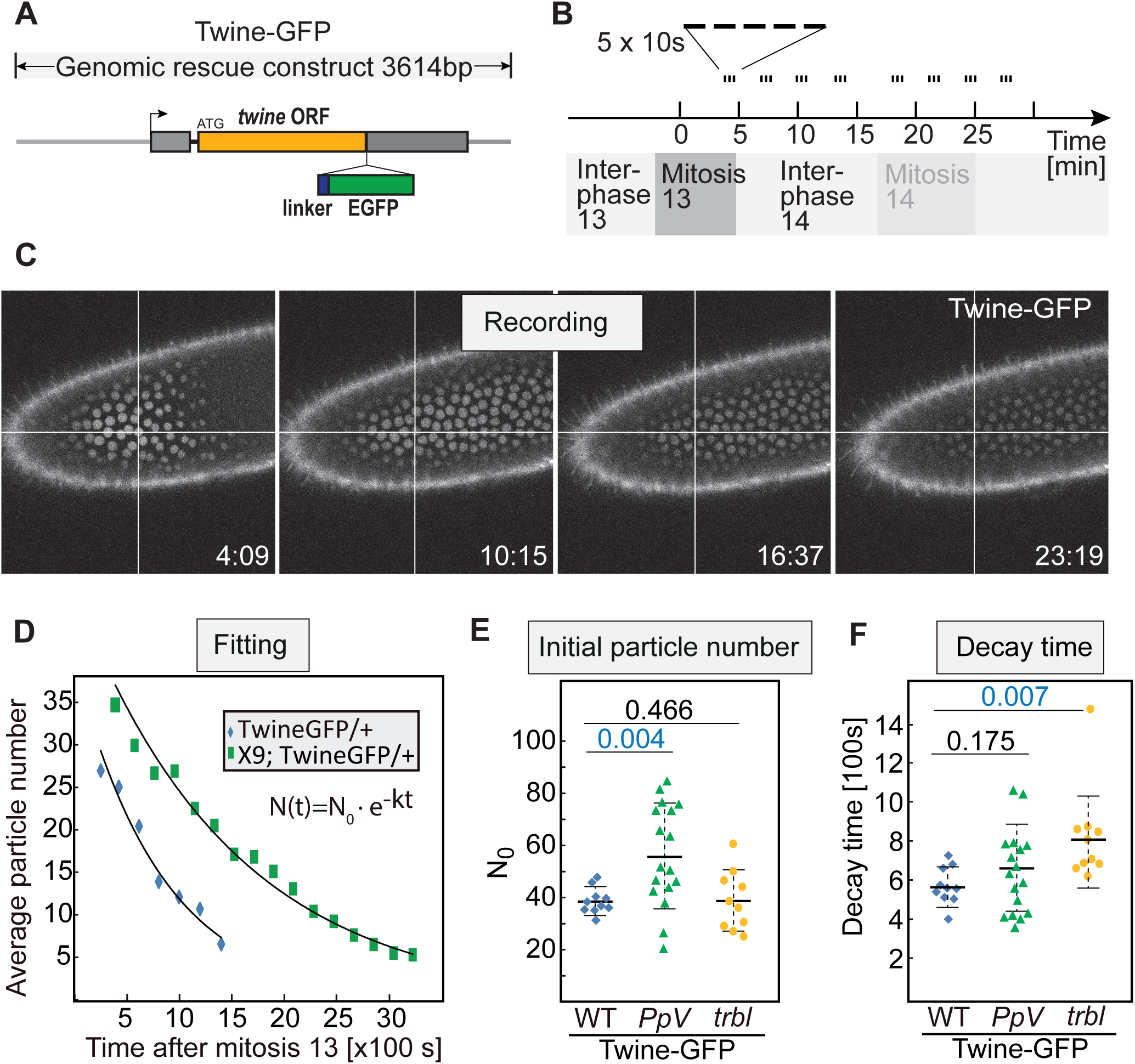
Expression profile of Twine-GFP. **(A)** Scheme of Twine-GFP genomic rescue construct. *twine* open reading frame (ORF), sindicated in orange. Linker sequence (Stuffer) is indicated in blue box and EGFP is in green box. Untranslated regions (UTR) are indicated in grey box. 3614 bp genomic region is depicted in grey line. **(B)** Experimental scheme. Five 10 s measurements are recorded for each time point (every 3 min). Multiple time points in interphase 14/15 are recorded for each embryo. **(C)** Images from a recording. **(D)** Representative time courses for a wild type and mutant embryo. Exponential fitting provides two parameters for each embryo, initial particle number (No) and the decay time (t). **(E, F)** Distribution of calculated initial particle number and decay time for wild type embryos, embryos from *P_p_V*[X9] germ line clones, and maternal and zygotic homozygous *trbl* embryos, all of which are with one copy of Twine-GFP. Me an, bold line, standard deviation, dashed line. The numbers indicate the statistical significance (p value) for the difference between two distributions.

### Twine is hyperphosphorylated in *P_p_V* mutants

To assess a potential control of Twine protein stability by phosphorylation, we analyzed Twine protein by mass spectrometry. We isolated Twine-GFP with GFP-binder from staged wild type and *P_p_V* mutant embryos (Fig. 6A). The isolated amounts of Twine-GFP protein were visible with Coomassie blue staining (Fig. 6B). Isolated and digested bands were analyzed for their peptide sequences and for phosphorylated residues. In wild type and mutant embryos, we achieved coverage of 59.4% and 70%, respectively, and we detected peptides 1.8 fold more frequently in wild type than in mutants (Fig. 6C, Suppl. data Tab. S1). Two dual phosphorylation sites were found in both wild type and *P_p_V* embryos. Thr203 + Ser205 and Thr394+Ser396 (Fig. 6C, Tab. 1). Importantly, an additional three sites were identified in *P_p_V* mutants: Ser41, at least one residue in the region between 59 and 67, and a clustered site close to the C-terminus, Ser405 and Ser412 (Fig. 6C, Tab. 1). These sites are evolutionary not conserved (Suppl. Data Fig. S2). The mass spectrometric analysis indicates that Twine is phosphorylated at additional sites in *P_p_V* mutants and suggests that P_p_V may act on Twine either directly or indirectly via other protein phosphatase or kinase.

**Figure 6:**
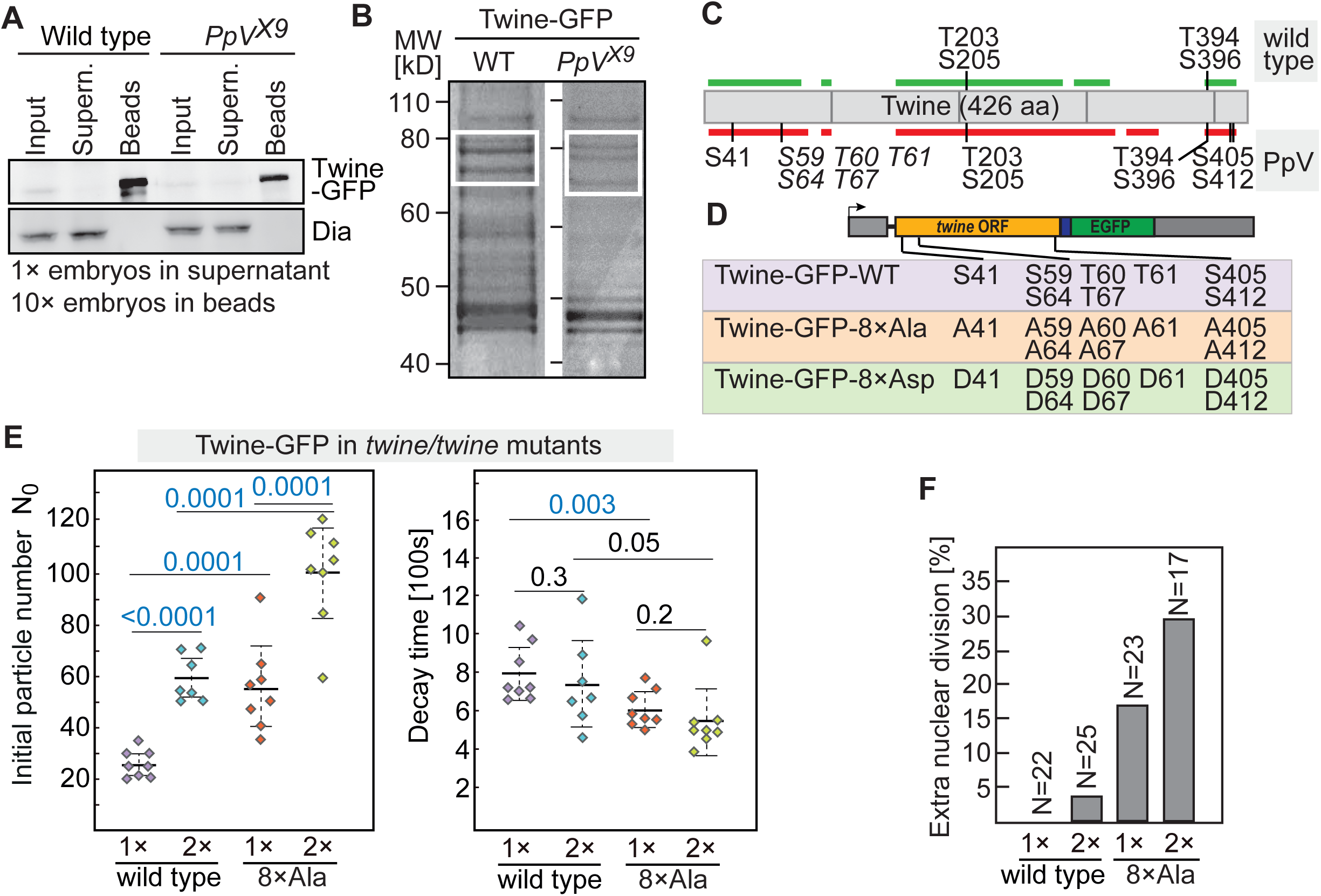
Identification of P_p_V dependent phosphorylation sites in Twine. **(A)** Twine-GFP was isolated with GFP-binder bound to beads from extracts of staged (0–1.5h) wild type embryos or embryos from *P_p_V*[X9] germ line clones. Western blot with GFP and Dia antibodies. **(B)** Image of Coomassie blue stained SDS polyacrylamide gel. White boxes indicate the area that was analyzed by mass-spectrometry. **(C)** Schematic diagram showing the position of Twine peptides covered in the analysis and the identified phosphorylation sites. italic types indicate ambiguous phosphosites. Coverage in wild type and *P_p_V*[X9] are shown in green and red, respectively. **(D)** Schematic diagram showing the transgenic Twine-GFP construct with phosphosites and phosphomimetic mutations. Eight P_p_V-dep endent residues are shown in the box of Twine-GFP-WT. Phosphosites mutant Twine-GFP-8×A la indicates the replacement of the eight phosphorylation residues to alanine/A. Phosphomimetic mutant Twine-GFP-8×As pindicates the replacement of the eight phosphorylation residues to aspartic acid /D. **(E)** Distribution of calculated initial particle number and decay time for embryos with one or two copies of wild type Twine-GFP and Twine-GFP-8×A la, all of which are in *twine* mutant background. Mean, bold line, standard deviation, dashed line. The numbers indicate the statistical significance (p value) for the difference between two distributions. **(F)** Frequency of additional nuclear division in the embryos of wild type Twine-GFP and Twine-GFP-8×Aia.

**Table 1.**
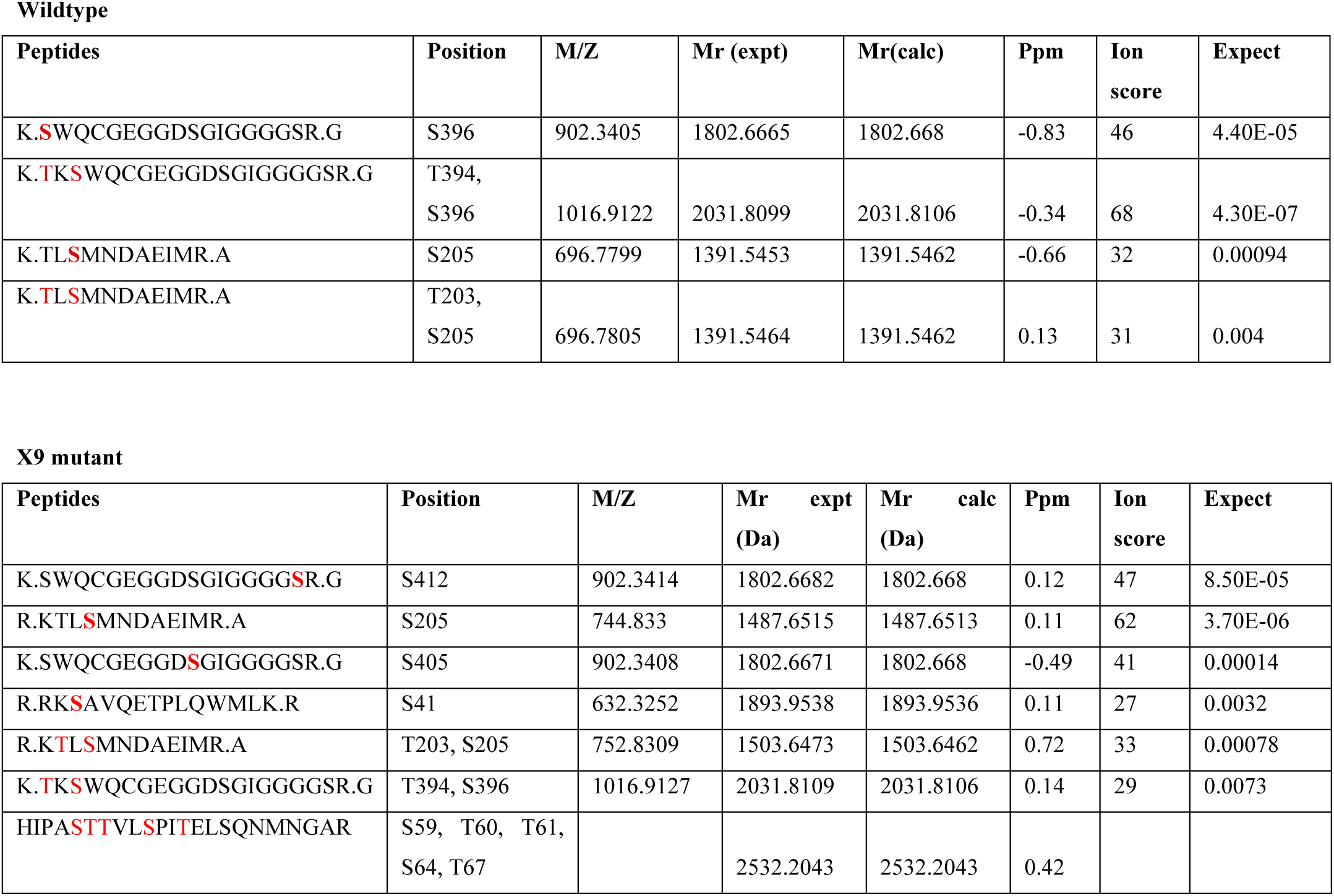
Analysis of phosphorylation sites by mass spectrometry. Results from Mascot data analysis of mass spectrometry (ms/ms) spectra. Peptide sequences are depicted of the Twine/Cdc25 as predict from fragmentation spectra. Predicted phosphorylation sites are marked in red (bold colors for unambiguous annotations). Although peptides were identified independently in many cases only highest scoring peptides are included. Observed mass/charge (M/Z) values indicate the result of the measurement and the calculated relative molecular weight (Mr) from the M/Z is indicated as experimental (expt) Mr in Dalton (Da). Mr calc depicts the calculated relative molecular weight in Dalton (Da) as calculated from the expected Mr from the database. Ppm indicates the error value between Mr expt and Mr calc and the ion score indicates the number of spectral ions matching the annotated fragments in the database. The expectation value is a statistical representation of the ion score expressed as p-value.

To test the relevance of the identified residues, we generated Twine-GFP constructs and corresponding transgenes, in which these residues were mutated (Fig. 6D). The transgenes were crossed into *twine* mutants to test for complementation as a sign for functionality of the mutated constructs. We generated phosphosite (Twine-GFP-8×Ala) and phosphomimetic (Twine-GFP-8×Asp) mutants in the context of a genomic rescue construct (Fig. 6D). Both mutated transgenes complemented the *twine* female sterility similar to the wild type transgene. However, the phosphomimetic mutant showed a dominant lethality with two copies. This dominant lethal phenotype indicates that Twine-GFP-8 ×Asp is acquired for new activities, such as new substrates. We did not further analyze the phosphomimetic mutant, as data interpretation would be questionable.

We measured Twine protein levels and their decay by fluorescence fluctuation analysis in embryos containing no endogenous Twine but the transgenic Twine-GFP, The phosphosite mutant contained a strongly increased amount of initial Twine levels (Fig. 6E, Tab. 2). We detected this increase in embryos with one copy of the transgene as well as in embryos with two copies. The decay rate was slightly lower in embryos with the phosphosite mutation than with the wild type allele (Fig. 6E, Tab. 2). Consistent with the higher Twine levels we observed a cell cycle phenotype. Amazingly, embryos with one or two copies of Twine-GFP-8×Ala frequently underwent an additional nuclear division (Fig. 6F). In summary, we detected developmentally relevant phosphorylation sites in Twine by our mass-spectrom etric analysis. Mutation of the P_p_V-dependent phosphorylation sites leads to higher protein expression and importantly a corresponding extra nuclear division in embryos comparable to *P_P_V* mutants. We conclude that P_p_V controls Twine phosphorylation state, ensures low expression levels and prompt decay of Twine, and thus timely remodeling of the cell cycle.

**Table 2.** Initial particle number and decay time measured by fluctuation analysis.

| Maternal genotype | No. of embryos | Initial particle number<br>Mean ± standard deviation | Decay time (100s)<br>Mean ± standard deviation |
| --- | --- | --- | --- |
| Twine-GFP/+ | 10 | 38.5 ± 5.0 (13%) | 571.7 ± 95.6 (17%) |
| *PpV*; Twine-GFP/+ (1) | 17 | 56.7 ± 19.5 (34%) | 640.1 ± 213.7 (33%) |
| *tribbles*, Twine-GFP/*tribbles* (2) | 10 | 38.9 ± 11.2 (29%) | 809.2 ± 256.1 (32%) |

| Genotype | No. of embryos | Initial particle number<br>Mean ± standard deviation | Decay time (100s)<br>Mean ± standard deviation |
| --- | --- | --- | --- |
| *twine*; Twine-GFP-WT/TM3 | 8 | 26.3 ± 4.7 (18%) | 799.9 ± 131.9 (16%) |
| *twine*; Twine-GFP-WT | 7 | 59.7 ± 8.7 (15%) | 736.0 ± 221.4 (30%) |
| *twine*; Twine-GFP-Ala/TM3 | 8 | 56.6 ± 16.2 (29%) | 601.9 ± 87.8 (15%) |
| *twine*; Twine-GFP-Ala | 8 | 99.9 ± 18.6 (19%) | 544.2 ± 173.2 (32%) |

(1) Females with PpV germline clones: *PpV*[X9] Frt18 hsFlp / ovoD Frt18 ; Twine-GFP / +
(2) Females with this genotype were crossed to *tribbles/tribbles* males

### *P_p_V* and *trbl* act in parallel

The differences in the dynamics of Twine-GFP suggest that P_p_V acts in a different manner on Twine than Trbl. To test this notion, we analyzed genetic interactions between *P_p_V* and *trb1*. *Trbl* has been implicated in cell cycle remodeling due to its zygotic expression and its ability to pause the nuclear division cycle (Grosshans 2000, Seher2000) and its role in degradation of Twine/Cdc25 (Farrell2013), *Trbl* homozygous animals have reduced viability, fertile flies, and *trbl* embryos from homozygous females undergo a normal number of nuclear divisions. In contrast, *trbl* females with *P_p_V* germ line clones did not produce any eggs. The ovaries of these females displayed multiple defects including egg chambers with an increased number of nurse cells (Data not shown). As both mutations are null alleles, we conclude that *P_p_V* and *trbl* act in parallel pathways in oogenesis, because the double mutant phenotype is stronger than the phenotypes of the single mutants.

In order to assess genetic interactions in embryos during MBT, we had to circumvent the oogenesis defect. We depleted *trbl* by injection of dsRNA into early embryos as described previously (Farrell2013). Following *trbl* RNAi injection in *P_p_V* mutant embryos we observed a slightly increased proportion of *P_p_V* embryos with an extra nuclear division cycle (Fig. 7B). Next, we tested whether the ability of *trbl* to induce a precocious cell cycle pause depended on *P_p_V.* Following injection of *trbl* mRNA into wild type as well as *P_p_V* embryos, we observed a precocious cell cycle pause as indicated by the lower nuclear density, larger nuclei at the injection site (Fig. 7A, 7B). These experiments show that *trbl* can precociously pause the cell cycle even in the absence of *P_p_V.* In summary, these experiments indicate that *P_p_V* and *trbl* act in separate pathways controlling entry into mitosis.

**Figure 7.**
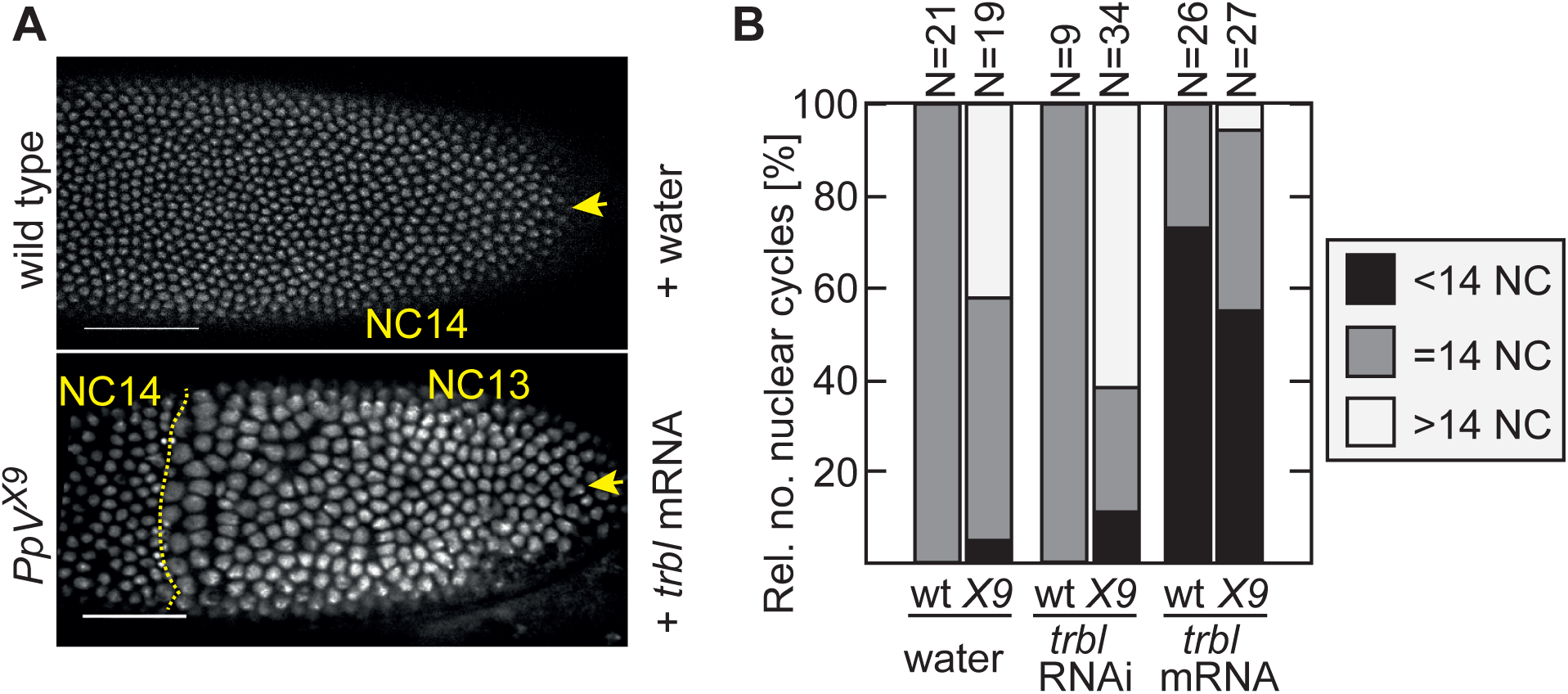
Functional interactions of *P_p_V* and *trbi*. Wild type embryos and embryos from germ line clones of *P_p_V*[X9] expressing Histone 2Av-RFP were injected with water, *trbi* mRNA, *trbi* ds RNA (RNAi) from the posterior pole (indicated by yellow arrow head). **(A)** Images from time lapse recordings of injected embryos. The number of nuclear cycle (NC) is indicated. Scale bar 50 μm, **(B)** Movies of injected embryos were scored for the number of nuclear divisions. Partial NC15 was scored as >14 NC, incomplete NC13 was <14 NC.

### The delayed cell cycle remodeling in *P_p_V* mutants does not depend on AuroraA

Next we investigated whether P_p_V controls Twine phosphorylation indirectly via AuroraA (AurA), which controls progression of mitotic events (Glover1995). It is assumed that PP-6 dependent dephosphorylation suppresses activation of AurA during mitosis in human cells (Zeng2010). AurA may also be a P_p_V substrate in Drosophila embryos. We observed an impaired chromosome segregation at low frequency in *P_p_V* mutants (NC10–13, 3% of the nuclei (N>100), three embryos) (Fig. 8A, Movie 2). Chromosomes did not separate and fused with chromosomes from neighboring spindles, although the arrangement of the centrosomes with single centrosomes at spindle poles appeared normal (Fig. 8A). In addition, we often observed delayed chromosome separation in telophase (Fig. 8B). In addition, we detected an additional band in an AurA western blot with extracts from *P_p_V* embryos in comparison to wild type extracts (Fig. 8C), which may correspond to a hyperphosphorylated AurA form. These observations are consistent with the antagonistic relationship of PP-6 and AurA in human cells.

**Figure 8.**
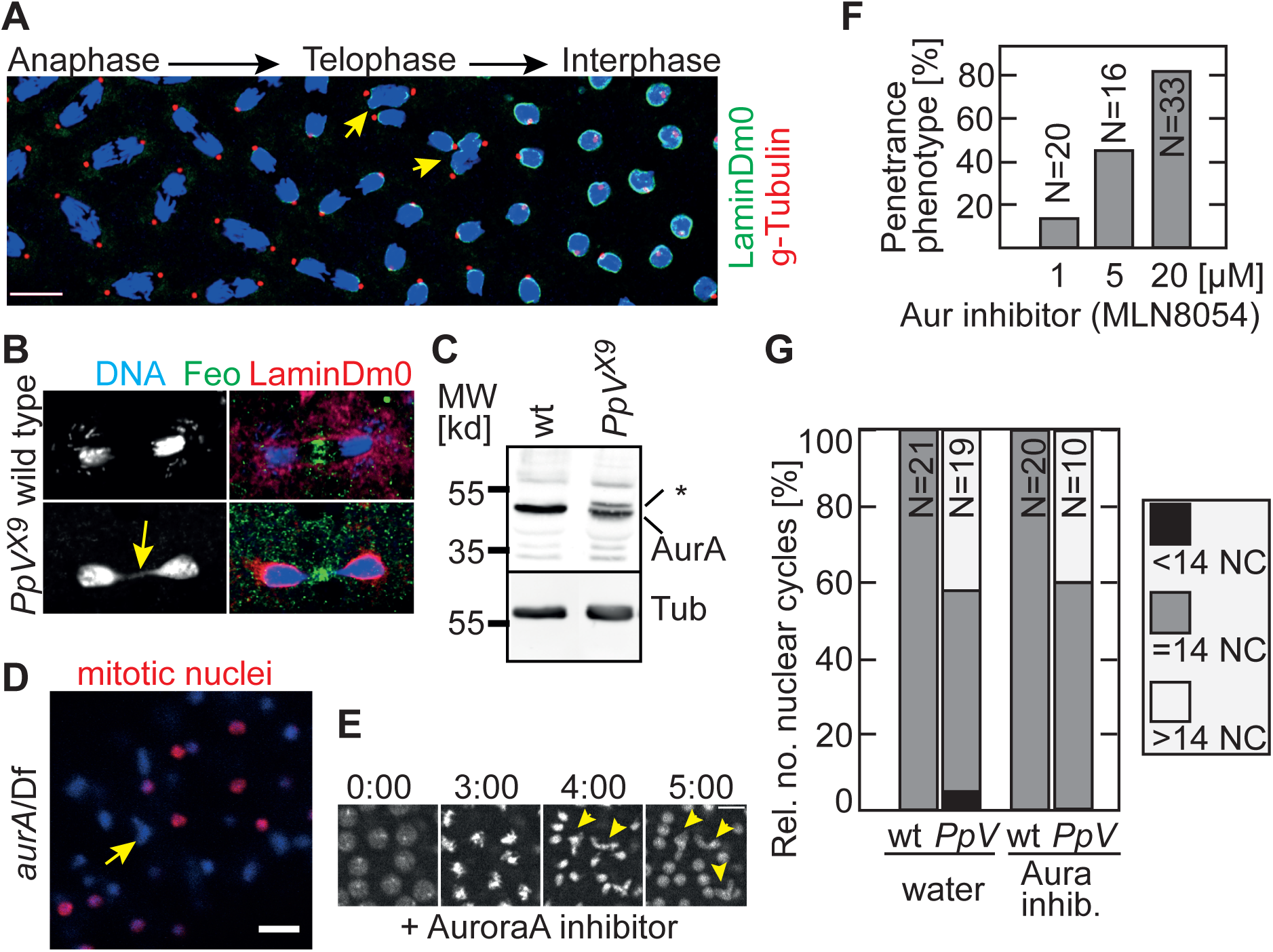
Relation of P_p_V and AuroraA. **(A)** Fixed embryos from *P_p_V*[X9] germ line clones were stained for centrosomes (γ-Tubulin, red), nuclear lamina (LaminDm0, green) and DNA (DAPI, blue). As a mitotic wave passed over the embryo, multiple mitotic stages were covered in a single embryo. Arrows in yellow point to mis-segregating nuclei. Scale bar 10 μm. **(B)** Fixed wild type embryo and embryo from *P_p_V*[X9] germ line clones stained nuclear lamina (LaminDmO, red), mid body (Feo, green) and DNA (DAPI, blue). The arrow in yellow points to delayed chromosome separation in telophase. **(C)** Western blot of wild type and *P_p_V*[X9] extracts with AurA and α-Tubulin antibodies. indicates an additional band in the P_p_V lane. **(D)** Image from fixed embryos from hemizygous *aurA* females stained for mitotic nuclei (p-histone3, red) and DNA (blue). Arrow in yellow points to nucleus with missegregated DNA. Scale bar 20 μm. **(E)** Images from time lapse recording of wild type embryos expressing histon 2Av-RFP injected with AuroraA inhibitor MLN8054 (20 μM). Scale bar 10 μm. Arrows in yellow point to missegregating spindles. **(F)** Dose dependence of mitotic phenotype following MLN8054 injection. **(G)** Movies of injected embryos were scored for the number of nuclear divisions. Partial NC15 was scored as >14 NC, incomplete NC13 was <14 NC.

It is conceivable that phosphorylated and thus activated AurA contributes to progression of nuclear division cycles. According to this model, AurA would have a higher activity in *P_p_V* mutants and thus trigger an extra nuclear division. A prediction from this model is that the extra division in *P_p_V* mutants depends on AurA. This prediction can be tested with double mutants of *P_p_V* and *aurA*. Embryos from hypomorphic *aurA* females develop until the blastoderm stage and are characterized by severe and asynchronous mitotic defects (Glover 1995, Fig. 8D), which prevented counting the number of nuclear divisions in *aurA* mutants and *P_p_V aurA* double mutants. To circumvent this problem, we employed the chemical inhibitor MLN8054, which has been demonstrated to be specific for AurA (Manfredi 2007). To validate the activity of this compound in Drosophila embryos, we injected MLN8054 into wild type embryos expressing Histone 2Av-RFP and recorded movies. Injected embryos frequently showed defects in chromosome segregation in anaphase and telophase of syncytial cycles depending on the concentration of the inhibitor (Fig. 8E). This phenotype is consistent with the *aurA* mutant phenotype (Glover 1995, Fig. 8D). The dose response curve indicates an effective concentration at about 20 μM in the injection solution (Fig. 8F). Injection into wild type embryos did not change the number of nuclear divisions on top of chromosomal segregation defects (Fig. 8G). All embryos went through 13 divisions. Injection into *P_p_V* mutant embryos resulted in a mixed phenotype with 13 or 14 nuclear divisions comparable to water injected *P_p_V* mutant embryos (Fig. 8G). We conclude that AurA activity is not required for the extra nuclear division in *P_p_V* mutant embryos. These data indicate that AurA does not link P_p_V to Twine for cell cycle remodeling during MBT.

## Discussion

Here we defined a novel function of P_p_V/PP-6 in cell cycle remodeling during MBT in *Drosophila.* At the center of this remodeling is the degradation of Twine/Cdc25 protein (Di Talia2013, Farrell2013). During the onset of cycle 14 and in response to MBT, the half life of Twine protein decreases from 20 min to 5 min (Di Talia2013). The biochemical signature of the degradation signal and the mechanism for the reduced half life remain elusive except for that Trbl and zygotic gene expression are involved. Here we revealed a mechanism that acts in addition to induced destabilization of Twine. The time, when Twine level falls below the critical threshold for entry into mitosis is determined by two parameters: (1) the decay time/half life and (2) initial levels. We propose a model, in which Trbl and other unknown zygotic factors controlled by the activated zygotic genome are involved in the induced destabilization during MBT, whereas maternal P_p_V controls the pre-MBT levels of Twine (Fig. 9A).

**Figure 9.**
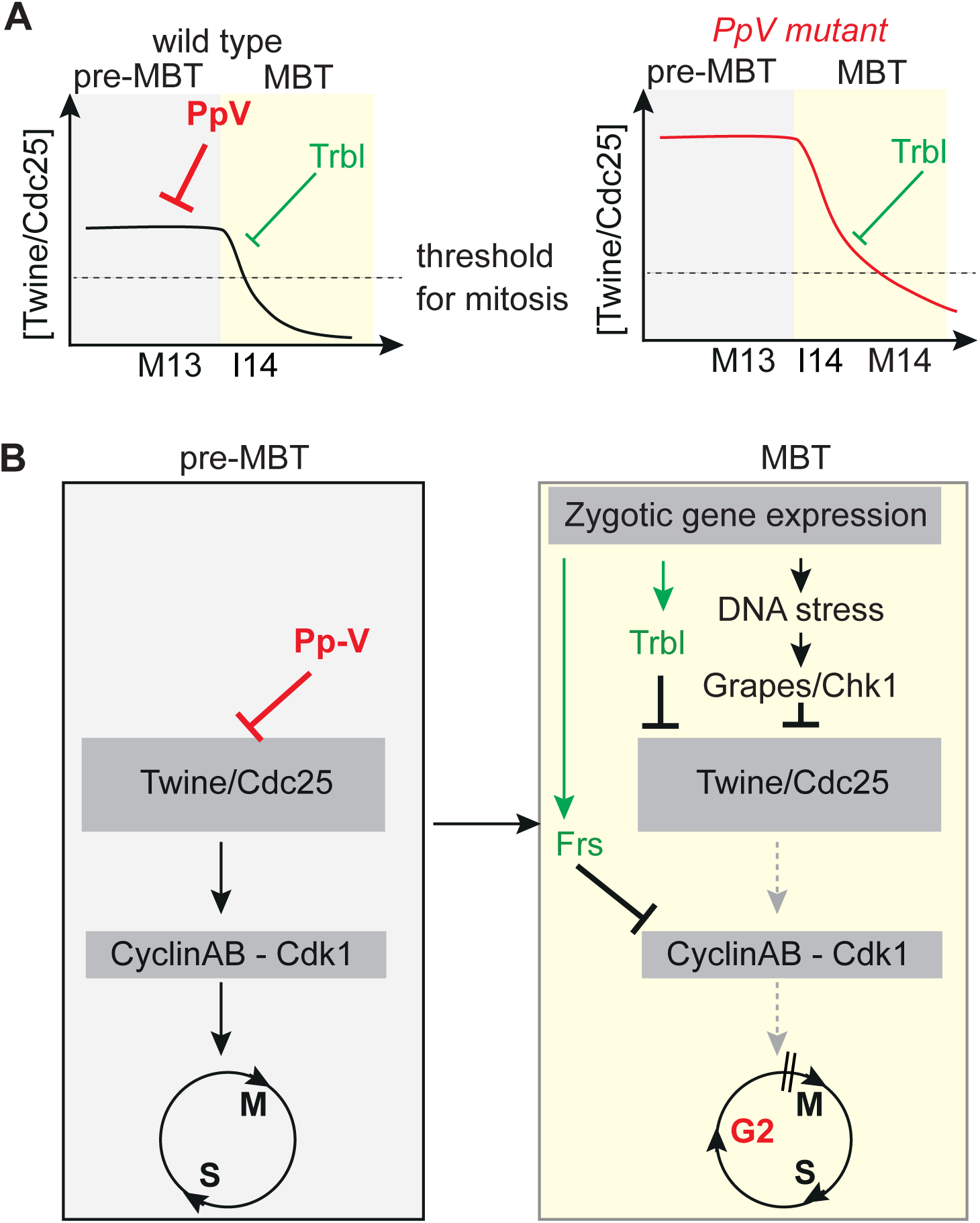
P_p_V function during MBT cell cycle control. **(A)** Schematic temporal profile of Twine/Cdc25 in wild type and *P_p_V* mutant embryos (red). *P_p_V* controls pre-MBT levels of Twine, whereas Trbl and other zygotic factors control the destabilization of Twine during MBT. **(B)** Cell cycle control during MBT. Activation of the zygotic genome leads to expression of the zygotic genes (*frs*, *trbl*) and DNA replication stress, which activates the DNA checkpoint. P_P_V constitutes a forth negative element in control Twine/Cdc25 and Cyclin-Cdk1.

Twine/Cdc25 protein is expressed in high levels during pre-MBT stage in many *P_p_V* mutant embryos indicating that P_p_V ensures low protein levels in wild type embryos. Beside the higher expression levels, it is remarkable that the embryo to embryo variation depends on P_p_V. In P_p_V mutants, we observed a wide variation of Twine. This variation is not due to limitations of our measurement technique, since measurements for nls-GFP and Twine-GFP in wild type embryos are robust with little variation. Our measurements indicate that P_p_V suppresses embryo to embryo variation by keeping Twine levels low. This would be consistent with a safeguarding function of P_p_V, which narrows the variation to a ground level. Such a mechanism could also explain the 30%–50% penetrance of *P_p_V* embryonic cell cycle phenotype.

P_p_V acts in parallel to other mechanisms ensuring the robust and timely switch in the cell cycle mode. In contrast to P_p_V, these are triggered by the onset of zygotic transcription either directly such as the mitotic inhibitor Frühstart (Frs) that targets the hydrophobic patch of the Cyclin-Cdk1 complex (Gawlinski2007) and Trbl that is involved in destabilization of Twine in MBT (Farrell2013), or indirectly such as the checkpoint kinase Grapes/Chk1 (Sung 2013, Blythe 2015b) (Fig. 9B).

Twine/Cdc25 may be a direct substrate, with P_p_V hydrolyzing the phosphates at the identified positions. Alternatively, P_p_V may dephosphorylate and thereby inhibit a protein kinase for the identified positions on Twine/Cdc25. As AuroraA is a target of P_p_V, P_p_V may act indirectly on Twine/Cdc25, in that P_p_V inactivates AuroraA, which then cannot phosphorylate Twine/Cdc25. Mitotic Cdc25b phosphorylation by AuroraA in cultured human tumor cells has previously been reported (Dutertre 2004). We do not favor this model, because firstly two of the three P_p_V dependent phosphorylation sites on Twine do not match the Aurora lenient consensus motif [KR].[ST][^^^P] (Sardon 2010). Secondly, AuroraA acts and is activated during mitosis, whereas control of Twine protein levels is an interphase process. Thirdly, and most importantly inhibition of AuroraA by chemical inhibitors in *P_p_V* mutants did not suppress the extra mitosis, indicating that the entry into an extra cycle does not depend on hyperactive AuroraA.

The cell cycle control during MBT is a remarkable robust process with almost no variation in wild type embryos. Such a robustness requires save guarding mechanisms against natural variation of protein content and egg size, for example, especially when thresholds and gradients (temporal decay of Twine) are involved. Multiple mechanisms, including control of Twine levels by P_p_V, have been revealed that control the remodeling from fast nuclear cycles to an embryonic cell cycle mode. Future experiments will define the mechanism how P_p_V controls Twine protein levels and many redundant pathways are needed for the robust cell cycle switch during MBT.

## Methods and materials

### Genetics

Fly stocks were obtained from the Bloomington Drosophila Stock Center, if not otherwise noted. Following fly strains and mutations were used. *trbl*[EP1119], *twine* [HB5]. Following transgenes were used. Twine-GFP (integrated at the landing site att P2/68A4, S. Blythe, S. Di Talia and E. Wieschaus), Histone2Av-GFP/RFP. Genomic transgenes (P_p_V[ +], GFP-P_p_V[ +]) were generated according to standard protocols by PhiC31 integrase-mediated site-specific insertions in the landing site ZH-86Fb (Bischof2007). The germ line clone phenotype on the X9 chromosome was mapped to the X:4–6 region and separated from another lethal mutation in the X:13–16 region by meiotic recombination with marker chromosomes. The lethality was mapped by complementation with duplications and deficiencies to 5F3–4 region delimited by the proximal breakpoint of Df(1)ED6829 and distal breakpoint of Dp(1;3)DC156. The duplication Dp(1;3)DC155 rescued lethality and the germ line clone phenotype. The genes in this region, i. e. *swaPsi*, *swa*, *Marf* and *P_p_V*, were sequenced on the X9 chromosome in comparison to the X238 chromosome, which was isolated in the same mutagenesis background.

### Molecular genetics, cloning, constructs

For the *P_p_V* rescue construct, a 2679bp EcoRI fragment from BAC clone 18C-18 (BACPAC Resources Center) was isolated and cloned into the pattB vector (Bischof2007). For GFP-P_p_V **the 5’ terminal 444 bp (a HindII/EcoRI**-BspM1 fragment) were replaced by a corresponding 1229 bp fragment with codon optimized GFP and a linker inserted at the start codon, which was synthesized by Eurofins Genomics and cloned into the pattB vector. CS2-tribbles plasmid template was linearized by XhoI and transcribed by SP6 Transcription Kit (Ambion). dsRNAs were produced by T7 RNA polymerase (Ambion), NTPs, RNase inhibitor (Thermo Fisher) and Pyrophosphatase (Thermo Fisher), using CS2-tribbles as template and ds RNA primers BL10 (GTAATACGACTCACTATAGGGCGATCAGCGCACAGCCTAGTCA) and BL11 (GTAATACGACTCACTATAGGGCGATGGCCATAGATGGTGCTCC). The Twine-EGFP transgene was synthesizing as a 3.6 kb complementing genomic fragment (Alphey1992) with an in-frame insertion of a Drosophila codon optimized EGFP at the C-terminus including a linker sequence (S. Blythe, S. Di Talia and E. Wieschaus). The constructs of mutated Twine phosphosites were synthesized by Eurofins Genomics and cloned into the KpnI/BamHI fragment of Twine-EGFP-pBABR plasmid.

### RNA isolation, quantitative PCR

Quantitative PCR and data analysis were carried out according to the protocols of SYBR Green Real-Time PCR Master Mixes and qPCR system software (Thermo Fisher Scientific). cDNA template was synthesized by reverse transcription of Drosophila total RNA, which was isolated by using TRIzol total RNA isolation protocol (Invitrogen). The following primer pairs were used: ***twine* qPCR primers BL16 (GAGTTCCTTGGCGGACACAT) and BL17 (CAGGATAGTCCAGTGCCGGAT); *GAPDH* qPCR primers MP37F (CACCAGTTCATTCCCAACTT) and MP37R (CTTGCCTTCAGGTGACGC).**

### Antibodies, Immunisation

The *P_p_V* coding sequence was cloned into an expression vector with a C-terminal His-tag. The P_p_V-His protein was purified under denaturing conditions (Trenzyme, Konstanz). Rabbits were immunized with the purified denatured protein (BioGenes, Berlin). In western blots the serum detected a band that was not present in extracts from *P_p_V*[X9] embryos. In whole mount staining no difference between wild type and *P_p_V*[X9] embryos was observed. Following antibodies were used: AuroraA (Giet 2002), Feo (rabbit, Verni2004), LaminDm0 (mouse, T47/1/1, Risau1981), γ-tubulin (GTU-88, Sigma), GFP-booster (Chromotek), Dia (rabbit, guinea pig, Grosshans 2005, Wenzl2010), Slam (rabbit, guinea pig, Acharya 2014), Twine (rat, Di Talia2013).

### Immunostaining

Formaldehyde or heat fixed embryos were rinsed thrice in PBT, and blocked in PBT with 5% BSA at 4 °C overnight. The primary antibodies were added in the respective dilutions in 0.1% BSA with PBT and embryos were incubated 2 h with constant rotation at room temperature. Then the embryos were rinsed thrice and washed four times 15 min with PBT. Secondary antibodies were added in PBT and embryos were incubated for 2 h. Embryos were rinsed thrice and washed four times 15 min in PBT again. Embryos were then stained with DNA dye, rinsed thrice in PBT, washed in PBT for 5 min and mounted in Aquapolymount (Polysciences).

### Embryo microinjection

Embryos were dechorionated with 50% Klorix bleach for 90 s, dried in a desiccation chamber for 5 to 10 min, then covered by halocarbon oil. Glass capillaries with internal filament were pulled as needles. For transgenesis, DNA was injected at 0.1 μg/μl posteriorly and prior to pole cell formation. Capped mRNAs were injected at a concentration of 800 ng/μl. dsRNA was injected at concentration of 12 mg/ml (Farrell 2013). MLN8054 Aurora Kinase inhibitor (Sellack chemicals, Manfredi 2007) was injected at a concentration of 20 μM diluted in water. We estimate the injection volume to be roughly 1–5% of the embryo volume.

### Western blot

Proteins were separated by SDS polyacrylamide gel electrophoresis and transferred to nitrocellulose membrane by semi-dry or wet transfer and stained with Ponceau S for loading control. Following blocking with 5% milk powder in PBT, the filters were incubated with primary antibodies in 0.5% BSA in PBT overnight at 4°C. After washing (3× rinsed, 4× 10 min in PBT) and incubation with secondary antibodies (800CW, 680CW, Donkey anti-guinea pig/mouse/rabbit IgG) for 1 h at room temperature, the blots were imaged with an Odyssey CLx Infrared imaging system. 16-bit images were processed by Photoshop and FIJI/Image J. Collected embryos (N=10-20) were pooled, dissolved in 1× Laemmli buffer and boiled at 95°C for 10 min. To facilitate homogenization, the lysates were gently triturated.

### Microscopy

Images of fixed and stained samples were recorded with a confocal microscope (Zeiss LS M780). The settings and objectives (25×/ water, 40×/ oil and 63 ×/water) were based on optimal imaging conditions and each channel was recorded separately. For life imaging, embryos were dechorionated with 50% hypochloride bleach for 90 s, aligned on a piece of apple-juice agar, transferred to a coated cover slip and covered with halocarbon oil. Live movies with differential interference contrast (DIC) optics were recorded with a light intensity of 2.5–3.0 V, an exposure time of 80–100 ms and a frame interval of 0.5–1 min. Fluorescent movies were recorded at an inverted spinning disc microscope (Zeiss, CSU-X1) with 30–50% laser intensity, 100 ms exposure time and a frame rate of 1 image per 0.5 to 1 min.

### Fluctuation analysis

Following female genotype were utilized: wild type 1×: Twine-GFP/+, *P_p_V* 1×: *P_p_V*[X9]/*ovoD*; Twine-GFP/+, *trbl* 1×: X9/*ovoD*; *trbl*, Twine-GFP/*trbl*. Phosphosite mutant 1×: *twine/twine*; Twine-GFP/TM3, *twine/twine*; Twine-GFP-8×Ala/TM3. Phosphosite mutant 2×: *twine/twine*; Twine-GFP/Twine-GFP, *twine/twine;* Twine-GFP-8×Ala/Twine-GFP-8×Ala. Fluctuation traces were recorded with a 63× oil immersion objective (Planapochromat, NA 1.4/oil) and a GaAsP detector on a confocal microscope (Zeiss LSM780) in fluorescence correlation spectrometry (FCS) mode. Onset of interphase 14 was defined as T=0. The data were analyzed as previously reported (Chen 2002, Digman 2008). Briefly, the intensity traces were analyzed using the number and brightness analysis. Time average <i> and variance **σ**^2^ were computed for each trace. From these values, we obtained the average brightness per molecule as <e> = (**σ**^2^/<i>) – 1 and the average number of molecules within the focal volume as <n> = <i>/<e>. The statistical significance (p value) of differences between the measured distributions were **calculated by Student–s t**-test. Usually 5 traces of each 10 s length were used for calculation of one data point N(t). F or control measurements with nls-GFP four data points were averaged. For the time dependent decay of Twine-GFP at least five data point were used for fitting in an exponential curve by linear regression analysis. Determination coefficients were in the range of >0.97.

### Mass-spec/phospho-sites

Dechorionated pre-syncytial and syncytial (0–1.5 h) Twine-GFP/Twine-GFP embryos and embryos from *P_p_V* germline clones with Twine-GFP (maternal genotype *P_pV_*[X9] F rt 18 hsFlp/ovoD Frt 18; Twine-GFP/+) were collected in large batches and snap frozen in liquid nitrogen. Each 10000 embryos were lysed with a dounce homogenizer in 1 ml lysis and washing buffer (50 mM Tris/HCl [pH 7.4], 500 mM NaCl, 1 mM DTT, 0.5 mM EDTA, 0.5% Tween 20, 1% Phosphatase Inhibitor Cocktail 3 (Sigma - Aldrich), 1 Tablet/50 ml Protease Inhibitor Cocktail (Roche)). The preparatory experiment was in the scale of about 50000 embryos. The embryonic lysate was centrifuged twice at 14,000 rpm at 4 °C for 15 min, and the supernatant was transferred into new Eppendorf tube. GFP-TrapA agarose beads (20 μl) were washed thrice with lysis and washing buffer and added to 1 ml of cleared embryonic lysate. After rotating on a wheel for 1 hour at 4 °C, spinning at 800 rpm for 2 min, and washing Twine-GFP thrice with lysis and washing buffer, protein was eluted from the beads in Laemmli buffer. Samples were run on SDS-PAGE 4–12% gradient 4–12% MOPS buffered system and bands excised and processed. Band slices were digested with trypsin over night at 30°C and the peptide extracted the next day. Samples were resuspended in 1% formic acid and a 15 μl Aiquots of each sample were run either before or after phosphopeptide enrichment. Phosphopeptides were enriched using Ti 4+ IMAC (ReSyn Biosciences) on an UltiMate 3000 RSLC nano system (Thermo Scientific) coupled to a LTQ OrbiTrap Velos Pro (Thermo Scientific). Peptides were initially trapped on an Acclaim PepMap 100 (C18, 100 μm × 2 cm) and then separated on an EasySpray PepMap RSLC C18 column (75 μm × 50 cm) (Thermo Scientific) over a 120 min linear gradient. The data was analyzed by Proteome Discoverer 1.4 (Thermo Scientific) using Mascot 2.4 (Matrix Science) as the search engine.

## Acknowledgements

We are grateful to H. Bastians, S. Blythe, G. Bucher, S. Di Talia, M. Gatti, S. Luschnig, H. Saumweber and E. Wieschaus for materials or discussions. We are grateful to M. Kojic, E. Özturk, S. Spangenberg, N. Vogt for help during preliminary experimental work. We thank R. Webster, D. Lamont and K. Beattie of the School of Life Sciences, University of Dundee, U.K. for sample processing and mass spectrometry. We acknowledge service support from the Developmental Studies Hybridoma Bank created by NICHD of the NIH/USA and maintained by the University of Iowa, the Bloomington Drosophila Stock Center (supported by NIH P40OD018537), the Drosophila Genomics and Genetic Resources, Kyoto, the BACPAC **Resources Center at Children’s Hospital Oakland and the Genomic Resource Center at Indiana** University (supported by NIH 2P40OD010949-10A1). BL was in part supported by a fellowship from the China Scholarship Council. HwS was in part supported by a predoctoral fellowship of the German Academic Exchange Service (DAAD). This work was in part supported by the Göttingen Centre for Molecular Biology (funds for equipment repair) and the Deutsche Forschungsgemeinschaft (DFG GR1945/3-1, SFB937/TP10 and equipment grant INST1525/16-1 FUGG).

## Author contributions

BL conducted the experiments and analyzed the data, HwS mapped and cloned the X9 mutation, IG analyzed the fluctuation data, HAM conducted the MS analysis, JG conceived the project, conducted the initial phenotypic analysis and wrote the manuscript.

## Conflict of interest

The authors have no conflict of interest.

## Supplemental data

**Figure S1.**
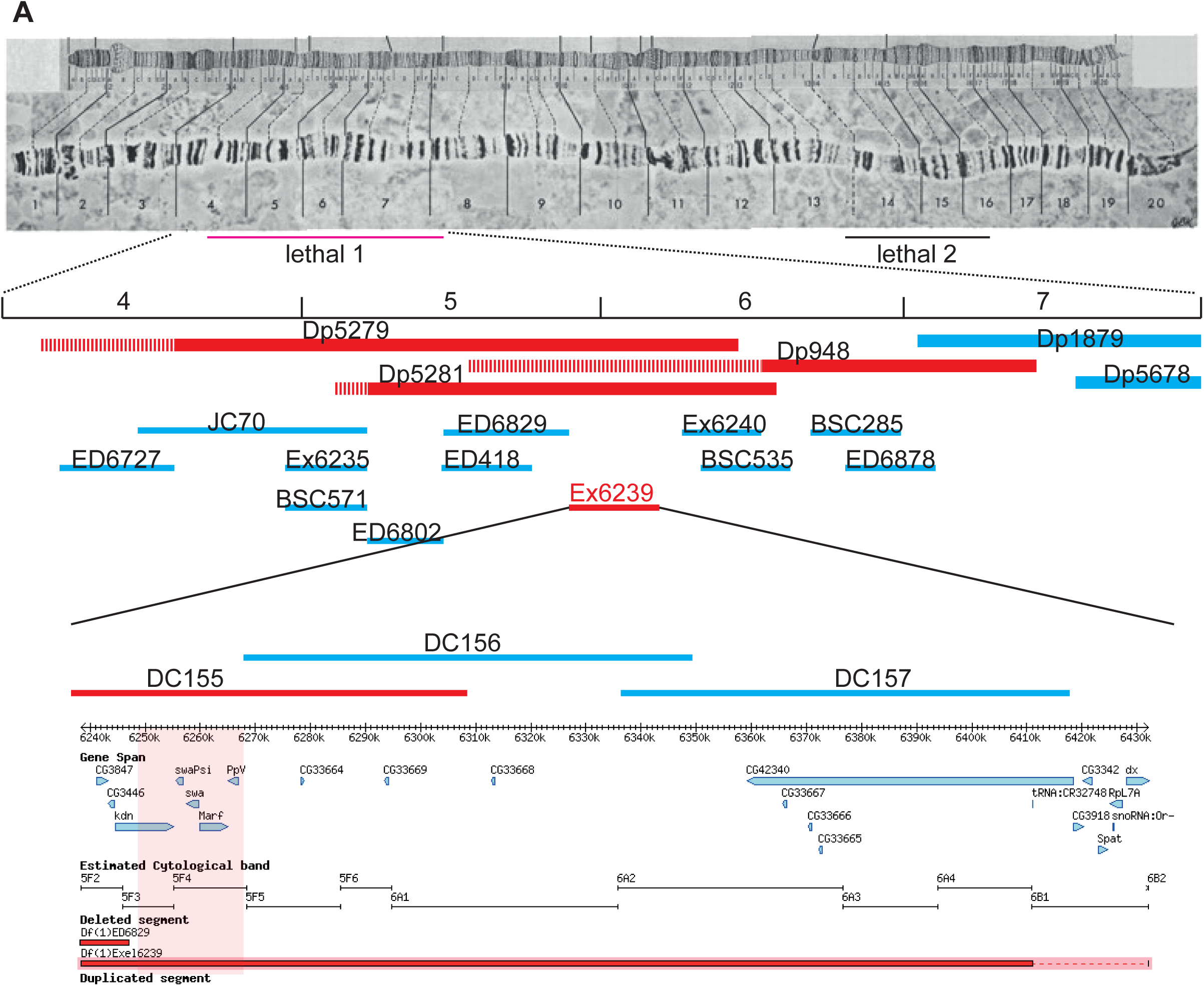
Mapping and cloning of X9. **(A)** Image of the X chromosome with the position of the mapped lethal mutations and mapped region by the closest breakpoints (distal Df(1)ED6829, proximal Dp(1;3)DC156). The distal lethality was mapped by complementation with indicated duplication and deficiency chromosomes. Dashed regions indicate uncertainty of the breakpoints. Complementing duplications and non-complementing deficiencies are marked in red. **(B)** Sequence traces of DNA amplified from heterozygous *P_p_V*[X9]/FM7 and in comparison X238/FM7 with a related genetic background. The double peak in the X9 trace reveals a non-sense mutation in codon 7 (TGG-> TAG).

**Figure S2.**
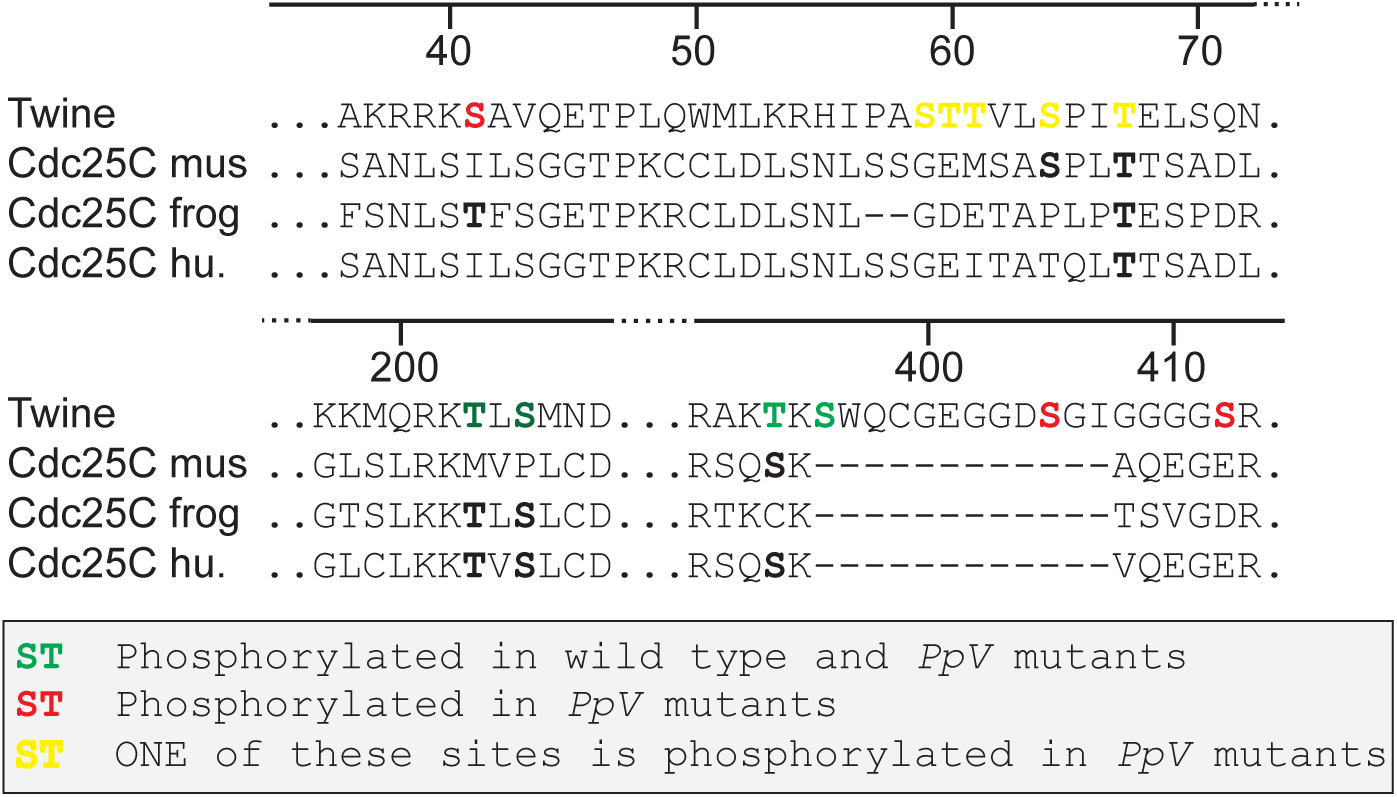
Alignment of Twine and Cdc25 homologue sequences. (mus-mouse, frog-Xenopus lav., hu-human). Phosphorylation sites are marked in color. Conserved residues are printed in bold.

**Table S1:** Comparison of non-phospho peptides in MS analysis of wild type and X9 mutant. The average peak area of five peptides were determined by XIC (extracted ion chromatograms) and compared between the wild type (wt) and the *P_p_V*[X9] mutant samples. The AA ratio of wt/X9 averages at 1.78 indicating that non-phosphopeptides were consistently about 1.8-fold more abundant in the wild type compared to *P_p_V*[X9] mutant samples.

| Peptide | mass | AA (wt) | AA (*PpV*) | AA ratio (wt/*PpV*) |
| --- | --- | --- | --- | --- |
| ALGDEPELIGDLSK | 728.8799 | 18358242 | 8486655 | 2.16 |
| LIQGEFDEQLGSQGGYEIDCR | 843.0633 | 2797579 | 1879327 | 1.49 |
| YPYEFLGGHIR | 676.3420 | 40374669 | 22168420 | 1.82 |
| IYVFHCEFSSER | 787.3584 | 8807155 | 5181341 | 1.7 |
| GQIQEAFPTLTSNQENR | 966.9741 | 44055348 | 25160476 | 1.75 |

**Movie 1** Movie of wild type embryo and embryo from *P_p_V*[X9] germ line clones expressing Histone2Av-RFP.

**Movie 2:** Movie of embryo from *P_p_V*[X9] germ line clones expressing Histone 2Av-RFP during mitosis.

## References

Acharya S, Laupsien P, Wenzl C, Yan S & Grosshans J (2014) Function and dynamics of slam in furrow formation in early Drosophila embryo. Dev. Biol. 386: 371–384

Afshar K, Werner ME, Tse YC, Glotzer M & Gönczy P (2010) Regulation of cortical contractility and spindle positioning by the protein phosphatase 6 PPH-6 in one-cell stage C. elegans embryos. Development 137: 237–247

Alphey L, Jimenez J, White-Cooper H, Dawson I, Nurse P & Glover DM (1992) twine, a cdc25 homolog that functions in the male and female germline of Drosophila. Cell 69: 977–988

Bastians H & Ponstingl H (1996) The novel human protein serine/threonine phosphatase 6 is a functional homologue of budding yeast Sit4p and fission yeast ppe1, which are involved in cell cycle regulation. J. Cell Sci. 109 (Pt 12): 2865–2874

Bischof J, Maeda RK, Hediger M, Karch F, Basler K (2007) An optimized transgenesis system for Drosophila using germ-line-specific Φ C31 integrases. Proc. Natl. Acad. Sci. U.S.A. 104: 3312-3317

Blythe SA & Wieschaus EF (2015a) Coordinating Cell Cycle Remodeling with Transcriptional Activation at the Drosophila MBT. Curr. Top. Dev. Biol. 113: 113–148

Blythe SA & Wieschaus EF (2015b) Zygotic Genome Activation Triggers the DNA Replication Checkpoint at the Midblastula Transition. Cell 160: 1169–1181

Chen F, Archambault V, Kar A, Lio P, D’Avino PP, Sinka R, Lilley K, Laue ED, Deak P, Capalbo L & Glover DM (2007) Multiple protein phosphatases are required for mitosis in Drosophila. Current Biology 17: 293–303

Chen Y, Müller JD, Ruan Q & Gratton E (2002) Molecular brightness characterization of EGFP in vivo by fluorescence fluctuation spectroscopy. Biophys. J. 82: 133–144

Di Talia S, She R, Blythe SA, Lu X, Zhang QF & Wieschaus EF (2013) Posttranslational control of Cdc25 degradation terminates Drosophila’s early cell-cycle program. Curr. Biol. 23: 127–132

Digman MA, Dalal R, Horwitz AF & Gratton E (2008) Mapping the number of molecules and brightness in the laser scanning microscope. Biophys. J. 94: 2320–2332

Dutertre S, Cazales M, Quaranta M, Froment C, Trabut V, Dozier C, Mirey G, Bouché J-P, Theis-Febvre N, Schmitt E, Monsarrat B, Prigent C & Ducommun B (2004) Phosphorylation of CDC25B by Aurora-A at the centrosome contributes to the G2-M transition. J. Cell Sci. 117: 2523–2531

Edgar BA & Datar SA (1996) Zygotic degradation of two maternal Cdc25 mRNAs terminates Drosophila’s early cell cycle program. Genes Dev. 10: 1966–1977

Ertych N, Stolz A, Valerius O, Braus GH & Bastians H (2016) CHK2-BRCA1 tumor-suppressor axis restrains oncogenic Aurora-A kinase to ensure proper mitotic microtubule assembly. Proc. Natl. Acad. Sci. U.S.A. 113: 1817–1822

Farrell JA & O’Farrell PH (2013) Mechanism and regulation of Cdc25/Twine protein destruction in embryonic cell-cycle remodeling. Curr. Biol. 23: 118–126

Farrell JA & O’Farrell PH (2014) From egg to gastrula: how the cell cycle is remodeled during the Drosophila mid-blastula transition. Annu. Rev. Genet. 48: 269–294

Foe VE, Odell GM & Edgar BA (1993) Mitosis and Morphogenesis in the Drosophila Embryo: Point and Counterpoint. In The Development of Drosophila melanogaster, Bate M & Arias AM (eds) Cold Spring Harbor Laboratory Press

Gawliński P, Nikolay R, Goursot C, Lawo S, Chaurasia B, Herz H - M, Kussler’Schneider Y, Ruppert T, Mayer M & Grosshans J (2007) The Drosophila mitotic inhibitor Frühstart specifically binds to the hydrophobic patch of cyclins. EMBO Rep. 8: 490–496

Giet R, McLean D, Descamps S, Lee MJ, Raff JW, Prigent C & Glover DM (2002) Drosophila Aurora A kinase is required to localize D-TACC to centrosomes and to regulate astral microtubules. J. Cell Biol. 156: 437–451

Glover DM, Leibowitz MH, McLean DA & Parry H (1995) Mutations in aurora prevent centrosome separation leading to the formation of monopolar spindles. Cell 81: 95–105

Grosshans J & Wieschaus E (2000) A genetic link between morphogenesis and cell division during formation of the ventral furrow in Drosophila. Cell 101: 523–531

Grosshans J, Müller HAJ & Wieschaus E (2003) Control of cleavage cycles in Drosophila embryos by frühstart. Dev. Cell 5: 285–294

Grosshans J, Wenzl C, Herz HM, Bartoszewski S, Schnorrer F, Vogt N, Schwarz H, Muller HA (2005) RhoGEF2 and the formin Dia control the formation of the furrow canal by directed actin assembly during Drosophila cellularisation. Development 132: 1009-1020

Hammond D, Zeng K, Espert A, Bastos RN, Baron RD, Gruneberg U & Barr FA (2013) Melanoma-associated mutations in protein phosphatase 6 cause chromosome instability and DNA damage owing to dysregulated Aurora-A. J. Cell Sci. 126: 3429–3440

Harrison MM & Eisen MB (2015) Transcriptional Activation of the Zygotic Genome in Drosophila. Curr. Top. Dev. Biol. 113: 85–112

Hodis E, Watson IR, Kryukov GV, Arold ST, Imielinski M, Theurillat J-P, Nickerson E, Auclair D, Li L, Place C, Dicara D, Ramos AH, Lawrence MS, Cibulskis K, Sivachenko A, Voet D, Saksena G, Stransky N, Onofrio RC, Winckler W, et al (2012) A landscape of driver mutations in melanoma. Cell 150: 251–263

Hu M-W, Wang Z-B, Teng Y, Jiang Z-Z, Ma X-S, Hou N, Cheng X, Schatten H, Xu X, Yang X & Sun Q-Y (2015) Loss of protein phosphatase 6 in oocytes causes failure of meiosis II exit and impaired female fertility. J. Cell Sci. 128: 3769–3780

Krauthammer M, Kong Y, Ha BH, Evans P, Bacchiocchi A, McCusker JP, Cheng E, Davis MJ, Goh G, Choi M, Ariyan S, Narayan D, Dutton-Regester K, Capatana A, Holman EC, Bosenberg M, Sznol M, Kluger HM, Brash DE, Stern DF, et al (2012) Exome sequencing identifies recurrent somatic RAC1 mutations in melanoma. Nat. Genet. 44: 1006–1014

Laver JD, Marsolais AJ, Smibert CA & Lipshitz HD (2015) Regulation and function of maternal gene products during the maternal-to-zygotic transition in Drosophila. Curr. Top. Dev. Biol. 113: 43–84

Luschnig S, Moussian B, Krauss J, Desjeux I, Perkovic J & Nüsslein-Volhard C (2004) An F1 genetic screen for maternal-effect mutations affecting embryonic pattern formation in Drosophila melanogaster. Genetics 167: 325–342

Liu B & Grosshans J (2017) Link of zygotic genome activation and cell cycle control. Methods Mol. Biol. 1605: 11–30

Manfredi MG, Ecsedy JA, Meetze KA, Balani SK, Burenkova O, Chen W, Galvin KM, Hoar KM, Huck JJ, LeRoy PJ, Ray ET, Sells TB, Stringer B, Stroud SG, Vos TJ, Weatherhead GS, Wysong DR, Zhang M, Bolen JB & Claiborne CF (2007) Antitumor activity of MLN8054, an orally active small-molecule inhibitor of Aurora A kinase. Proc. Natl. Acad. Sci. U.S.A. 104: 4106–4111

Mann DJ, Dombradi V & Cohen PTW (1993) Drosophila protein phosphatase. 12: 4833–4842

Ogoh H, Tanuma N, Matsui Y, Hayakawa N, Inagaki A, Sumiyoshi M, Momoi Y, Kishimoto A, Suzuki M, Sasaki N, Ohuchi T, Nomura M, Teruya Y, Yasuda K, Watanabe T & Shima H (2016) The protein phosphatase 6 catalytic subunit (Ppp6c) is indispensable for proper post-implantation embryogenesis. Mech. Dev. 139: 1–9

Risau W, Saumweber H & Symmons P (1981) Monoclonal antibodies against a nuclear membrane protein of Drosophila. Localization by indirect immunofluorescence and detection of antigen using a new protein blotting procedure. Exp. Cell Res. 133: 47–54

Rusin SF, Schlosser KA, Adamo ME & Kettenbach AN (2015) Quantitative phosphoproteomics reveals new roles for the protein phosphatase PP6 in mitotic cells. Sci Signal 8: rs12–rs12

Sardon T, Pache RA, Stein A, Molina H, Vernos I & Aloy P (2010) Uncovering new substrates for Aurora A kinase. EMBO Rep. 11: 977–984

Seher TC & Leptin M (2000) Tribbles, a cell-cycle brake that coordinates proliferation and morphogenesis during Drosophila gastrulation. Current Biology 10: 623–629

Stefansson B & Brautigan DL (2007) Protein phosphatase PP6 N terminal domain restricts G1 to S phase progression in human cancer cells. Cell Cycle 6: 1386–1392

Sung H-W, Spangenberg S, Vogt N & Grosshans J (2013) Number of nuclear divisions in the Drosophila blastoderm controlled by onset of zygotic transcription. Curr. Biol. 23: 133–138

Verni F, Somma MP, Gunsalus KC, Bonaccorsi S, Belloni G, Goldberg ML & Gatti M (2004) Feo, the Drosophila homolog of PRC1, is required for central-spindle formation and cytokinesis. Curr Biol 14: 1569–1575

Vogt N, Koch I, Schwarz H, Schnorrer F & Nüsslein-Volhard C (2006) The gammaTuRC components Grip75 and Grip128 have an essential microtubule-anchoring function in the Drosophila germline. Development 133: 3963–3972

Wenzl C, Yan S, Laupsien P & Grosshans J (2010) Localization of RhoGEF2 during Drosophila cellularization is developmentally controlled by slam. Mech. Dev. 127: 371–384

Yan S, Xu Z, Lou F, Zhang L, Ke F, Bai J, Liu Z, Liu J, Wang H, Zhu H, Sun Y, Cai W, Gao Y, Su B, Li Q, Yang X, Yu J, Lai Y, Yu X-Z, Zheng Y, et al (2015) NF-κB-induced microRNA-31 promotes epidermal hyperplasia by repressing protein phosphatase 6 in psoriasis. Nat Commun 6: 7652

Zeng K, Bastos RN, Barr FA & Gruneberg U (2010) Protein phosphatase 6 regulates mitotic spindle formation by controlling the T-loop phosphorylation state of Aurora A bound to its activator TPX2. J. Cell Biol. 191: 1315–1332

Zhong J, Liao J, Liu X, Wang P, Liu J, Hou W, Zhu B, Yao L, Wang J, Li J, Stark JM, Xie Y & Xu X (2011) Protein phosphatase PP6 is required for homology-directed repair of DNA double-strand breaks. Cell Cycle 10: 1411–1419

